# Phenotypic plasticity in chemical defence allows butterflies to diversify host use strategies

**DOI:** 10.1101/2020.04.07.030122

**Authors:** Érika C. P. de Castro, Jamie Musgrove, Søren Bak, W. Owen McMillan, Chris D. Jiggins

## Abstract

Hostplant specialization is a major force driving ecological niche partitioning and diversification in insect herbivores. The cyanogenic defences of *Passiflora* plants keeps most herbivores at bay, but not larvae of *Heliconiu*s butterflies, which can both sequester and biosynthesize cyanogenic compounds. Here, we demonstrate that both *Heliconius cydno chioneus*, a host plant generalist, and *H. melpomene rosina, a specialist,* have remarkable plasticity in their chemical defence. When feeding on *Passiflora* species with cyanogenic compounds they can readily sequester, both species downregulate the biosynthesis of these compounds. In contrast, when fed on *Passiflora* plants that do not contain cyanogenic glucosides that can be sequestered, both species increase biosynthesis. This biochemical plasticity comes at a significant fitness cost for specialist like *H. m. rosina*, as growth rates for this species negatively correlate with biosynthesis levels, but not for a generalist like *H. c. chioneus*. In exchange, *H. m rosina* has increased performance when sequestration is possible as on its specialised hostplant. In summary, phenotypic plasticity in biochemical responses to different host plants offers these butterflies the ability to widen their range of potential host within the *Passiflora* genus, while maintaining their chemical defences.

## INTRODUCTION

Phenotypic plasticity is widely recognised as an adaptation that allows organisms to survive in a variable environment [1]. Furthermore, there is interest in the idea that plasticity might permit populations to invade otherwise inaccessible niches or habitats [2][3]. Hostplant specialization is undoubtedly one of the most important forces driving diversification and shaping niche dimension for phytophagous insects [4][5]. Specialized insects often evolved not only to handle the chemical defences of their favorite hosts, but also to become dependent on plant compounds [6]. Whereas inducible defences of plants by herbivory has been well studied [7][8][9][10], there has been relatively little exploration of the mechanisms of biochemical plasticity in insect herbivores that could allow them to exploit diverse hosts [11].

The vast majority of aposematic butterflies acquired their toxic compounds from their larval hosts through sequestration. For example, the monarch butterfly sequesters cardenolides from milkweeds; swallowtails obtain Aristolochic acids from Aristolochiaceae; Ithomiini sequester pyrrolizidine alkaloids mostly from Solanaceae; and some toxic lycaenids acquired cycasin from Cycadales [6]. Sequestration of plant toxins during larval feeding is an adaptation that arose in many butterfly groups and plays an important role in the antagonist coevolution with their hosts. In contrast to most butterflies, *Heliconius* species have the ability to both sequester and synthesise their own chemical defences. All *Heliconius* butterflies can *de novo* biosynthesize aliphatic cyanogenic glucosides (CNglcs) using the amino acids valine and isoleucine as precursors [12] (see Figure 1 for CNglcs structures). Their *Passiflora* host plants are also chemically defended by a broad range of CNglcs [13], of which *Heliconius* can sequester aromatic, aliphatic, and especially simple cyclopentenyl during larval feeding [14][15][16]. To prevent sequestration, plants have responded by chemically modifying their defensive compounds. As an example, *H. melpomene* larvae can sequester cyclopentenyl CNglcs but cannot sequester sulfonated cyclopentenyl CNglcs from *P. caerulea* [15]. Other modified cyclopentenyl CNglcs, such the bis-glycosilated CNglcs, passibiflorin from *P. biflora*, have not yet been tested for sequestration. Disabling sequestration would not make these plants distasteful or toxic for *Heliconius*, but it could reduce their l value as a host and have deleterious effects on their fitness. From the perspective of the herbivores therefore, switching between biosynthesis and sequestration of toxins could allow butterflies to colonise a wider array of potential host plants independently of sequestration, while also maintaining their chemical defences. Here, we explore phenotypic plasticity in this trade-off in two *Heliconius* species with different host use strategies.

**Figure 1.**
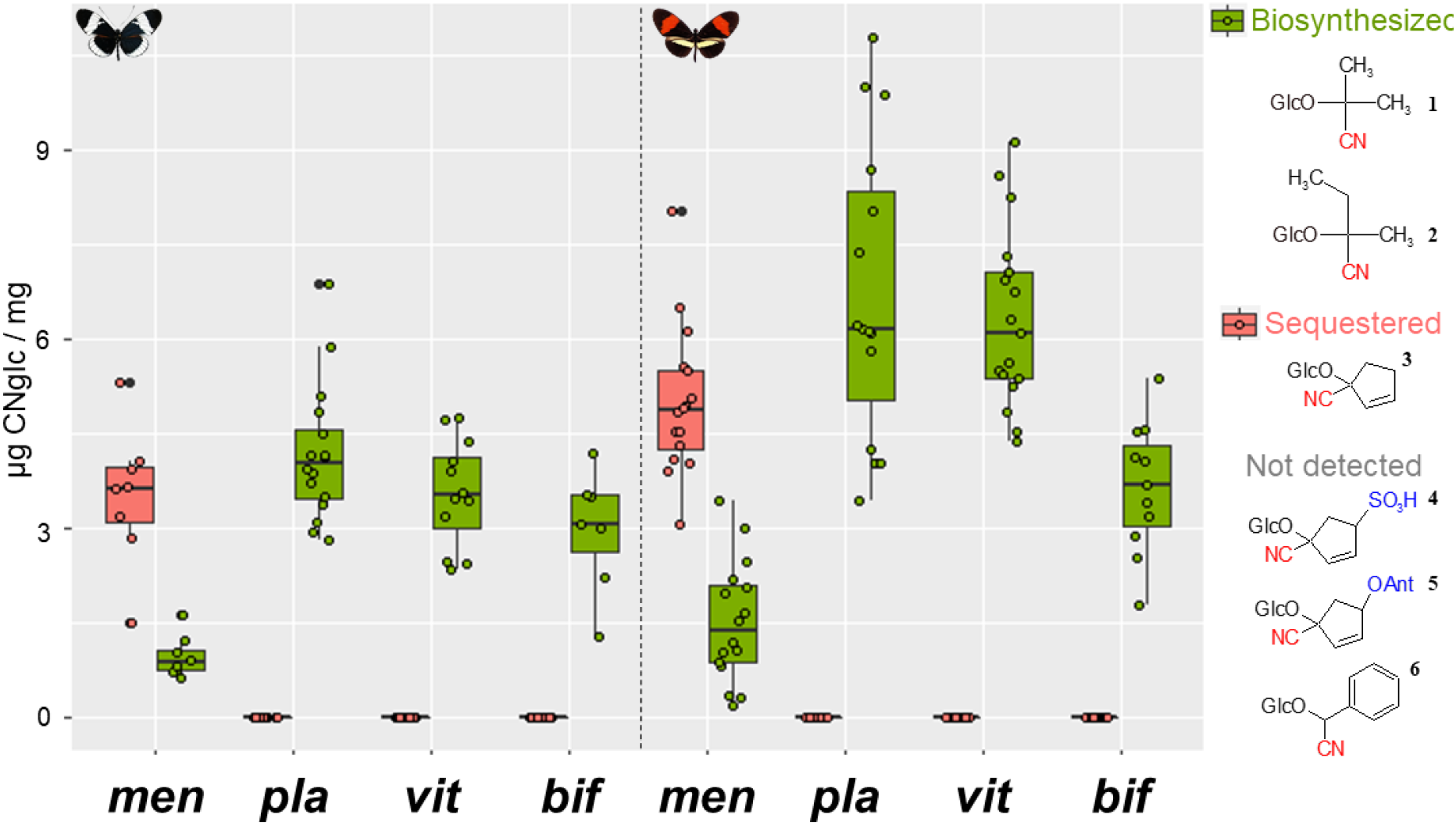
CNglc composition of *H. cydno* (left) and *H. melpomene* (right) raised on different *Passiflora* diet. Legend: vit= *P. vitifolia,* pla= *P. platyloba,* men= *P. menispermifolia*; bif= *P. biflora*. Green boxplots correspond to the biosynthesized cyanogens, linamarin^1^ and lotaustralin^2^, found in all butterflies. Salmon boxplots correspond to the sequestered CNglcs deidaclin^3^ only detected in butterflies raised on *P. menispermifolia*. Tetraphyllin B-sulphate^4^, passibiflorin^5^ and prunasin^6^ were not detected in butterflies, even thought they were present in the food plants *P. vitifolia*, *P. biflora* and *P. platyloba*, respectively. CNglcs present in each host plant is described in Table 1.

The closely related species *Heliconius melpomene* and *Heliconius cydno* (diverged ~2 MYA) are often found in sympatry and their reproductive isolation is not complete (hybrid males are fertile) [17]. *H. melpomene* is widespread in tropical America and lays eggs on several *Passiflora* species, but where it co-occurs with *Heliconius cydno*, is an ecological specialist, ovipositing mainly on *P. menispermifolia* (Panama) or *P. oerstedii* (Costa Rica and Colombia), although larvae are able to feed on a variety of species. In contrast, *H. cydno* is more generalist and oviposits on many *Passiflora* species [18][19][20][21]. These differences in oviposition preferences are genetically controlled [21]. Broadly, larval mortality and growth of both species are similar on different hosts [22], but a field experiment showed slightly higher establishment probability for *H. m. rosina* on *P. menispermifolia* [21]. Overall, experiments to date show only weak evidence for any adaptive advantage to the host specialisation of *H. melpomene* as compared to the more generalist strategy of *H. cydno*.

Nonetheless, these species show different host use strategies and feed on a variety of host plants with different chemical composition. Here, we take advantage of this ecology to explore plasticity in the balance between sequestration and biosynthesis of cyanogenic compounds among these two butterfly species, fed on four *Passiflora* species that produce different CNglcs (Table 1). We also examine growth rates to explore whether there are possible trade-offs in fitness when feeding on different host plants or adopting different strategies of chemical defence. Phenotypic plasticity in sequestration versus biosynthesis of CNglcs defences could facilitate host switching and diversification of *Heliconius* across the *Passiflora* radiation.

**Table 1.** The CNglcs composition of the *Passiflora* species utilized in this study.

|  | Aliphatic<br>CNgls | Aromatic<br>CNgls | Simple<br>Cyclopentenyl<br>CNgls | Modified<br>Cyclopentenyl<br>CNgls |
| --- | --- | --- | --- | --- |
| *P. vitifolia* | - | - | - | **Tetraphyllin B**<br>**sulphate** |
| *P. platyloba* | - | **Prunasin** | - | - |
| *P. menispermifolia* | - | - | **Deidaclin** | - |
| *P. biflora* | - | - | - | **Passibiforin** |

## METHODS

### Butterfly rearing

Butterflies used in this study were reared at the Smithsonian Tropical Research Institute, Gamboa, Panama. Mated female stocks of *H. cydno chioneus* and *H. melpomene rosina* were maintained in insectary cages and fed *ad libidum* with flowers (*Psiguria triphylla*, *Gurania eriantha*, *Psychotria poeppigiana*, *Lantana sp.*) and artificial nectar (10% sugar solution). Plants of one of the four species used in the experiment - *P. biflora, P. menispermifolia*, *P. platyloba*, and *P. vitifolia* - were always kept in cages for oviposition. Eggs were collected daily from host plants and kept in closed plastic cups) until hatching. On the morning of hatching, larvae were transferred to treatment-specific cages onto individual shoots. Young shoots with no evidence of herbivory were selected to minimise the effects of variable host quality. Suitable shoots were sterilized and placed into water-filled bottles sealed with cotton. Cages were checked every day and fresh shoots provided regularly. Pupae were immediately removed, weighed after one day of pupation and taped on the lid of individual 350 ml plastic tubes. After eclosion, butterflies were left in their individual tubes for a few hours to dry their wings and then removed for body measurements: total weight, forewing length and body length. Body length was measured from the end of the head to the end of the abdomen using mechanical callipers, and forewing length was measured from the central base to the most distal point. Butterflies were added into tubes containing 1.5 mL methanol 80% (v/v), sealed with Parafilm and stored at 4 °C.

### Chemical Analyses

Samples were homogenized in 1.5 methanol 80% (v/v) where they were soaked and centrifuged at 10,000 x g for 5 min. Supernatants were collected and kept in HPLC vials at −20 °C. Sample aliquots were filtered (Anapore 0.45 μm, Whatman) to remove insoluble components and diluted 50X times (v/v) in ultrapure water and injected into an Agilent 1100 Series LC (Agilent Technologies, Germany) hyphenated to a Bruker HCT-Ultra ion trap mass spectrometer (Bruker Daltonics, Bremen, Germany). Chromatographic separation was carried out using a Zorbax SB-C18 column (Agilent; 1.8μM, 2.1×50mm). MS and LC conditions are described in [16]. The sensitivity of the analytical system was monitored by running a pooled sample after each 20 experimental sample.

Sodium adducts of CNglcs detected in the butterflies were identified by comparing their m/z fragmentation patterns and RTs to authentic standards (Jaroszewski et al. 2002; Møller et al. 2016). Quantification of CNglcs present were estimated based on the Extracted Ion Chromatogram peak areas of each compounds and calculated from a standard curve of linamarin, lotaustralin, and amygdalin.

### Statistical Analyses

All statistical analyses were performed using R version 3.5.1 (R Core Team, 2017). Two-ways ANOVA was used to examine the interaction between butterfly species, larval diet, and sex for different biological traits (pupal weight, adult weight, forewing length, body size, and total CNglcs). One-way ANOVA followed by Tukey HSD tests were used to make pairwise comparisons between diets and analyse, within butterfly species, the effects of each diet on the measured traits.

## RESULTS

Larval diet affected the CNglc composition in adult butterflies of *H. melpomene* and *H. cydno* (Figure 1). Both species sequestered deidaclin when fed as larvae on *P. menispermifolia* plants, although *H. melpomene* sequestered significantly more deidaclin than *H. cydno* (ANOVA, *F*_1,22_= 8.851; *p*= 0.00699). Sequestration of deidaclin from *P. menispermifolia* was associated with a reduction of linamarin and lotaustralin biosynthesis in comparison with other diets. The modified CNglc passibiflorin from *P. biflora* and tetraphyllin B-sulphate from *P. vitifolia* were not found in both butterfly species raised on these diets, suggesting that they cannot sequester these compounds. Surprisingly, prunasin recently found in the haemolymph of larvae raised on *P. platyloba* [15] was not present in adults of either butterfly species. Instead, the derivative prunasin amide was found in adults reared on *P. platyloba* suggesting that they sequestered prunasin, but turned over into this compound to the corresponding amide during pupation.

Larval diet not only influenced the composition, but also the total concentration of CNglcs in both species (ANOVA, *H. cydno: F*_3,39_ = 3.653, *p*= 0.0205; *H. melpomene: F*_3,55_= 8.776, *p*= 0.00007) (Figure 2A). Both species had less CNglcs when reared on *P. biflora,* which they normally do not use as a host. On average, butterflies also had a higher CNglcs content when reared on *P. menispermifolia* than on *P. platyloba* and *P. vitifolia,* though these differences were only statistically significant for *H. cydno.* CNglc concentrations in *H. cydno* (3.85 ± 1.08) were on average lower than *H. melpomene* (5.96 ± 1.97).

**Figure 2.**
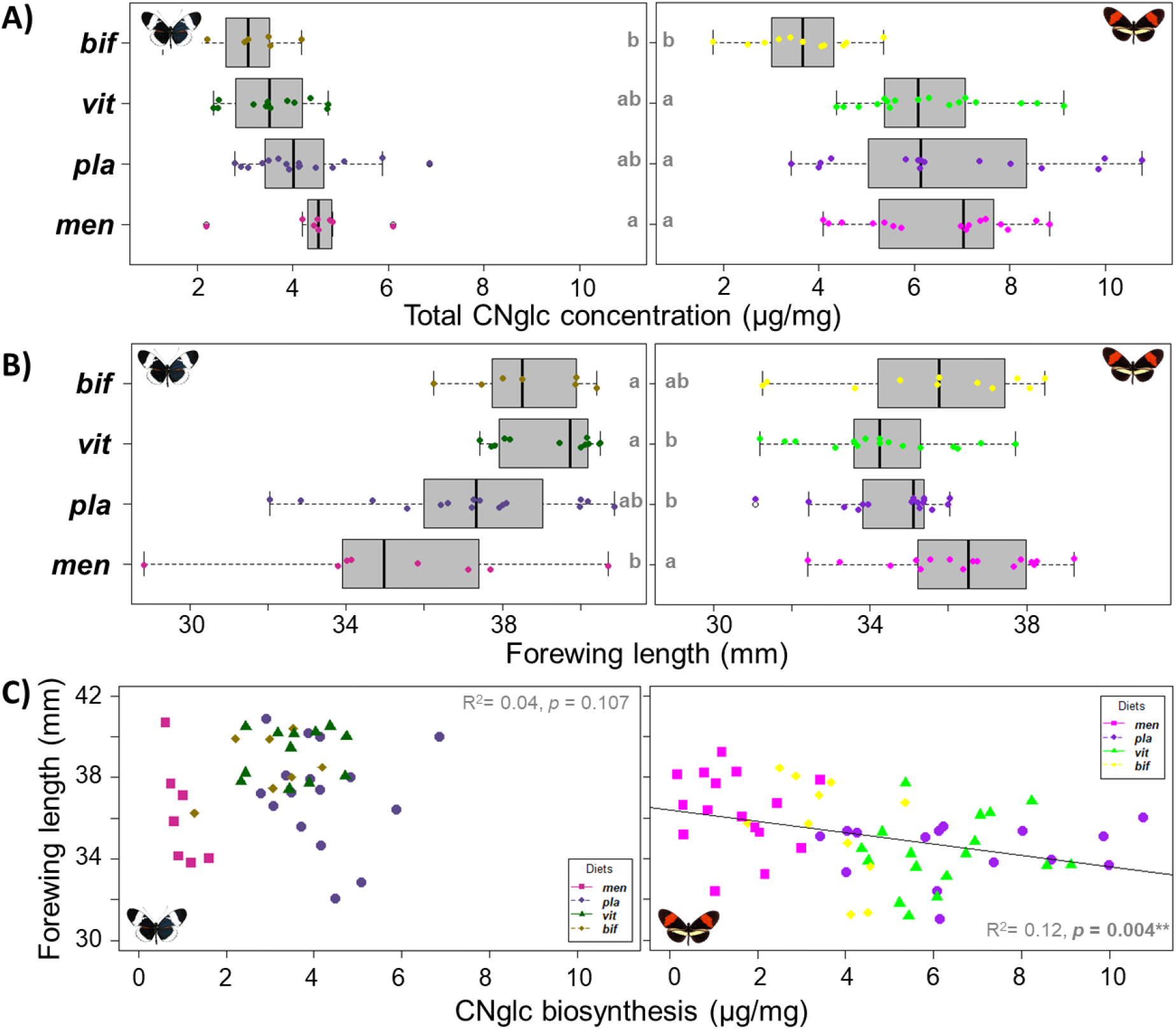
Effect of larval diet on **A)** total CNglcs concentration and **B)** forewing length of *H. cydno* (left) and *H. melpomene* (right). Letters over boxplots correspond to post-hoc comparisons (Tukey HSD) within butterfly species, where different letters indicate statistically significant treatments. C) Correlation between forewing length and concentration of biosynthesized CNglcs in *H. cydno* (left) and *H. melpomene* (right). Legend: vit= *P. vitifolia,* pla= *P. platyloba,* men= *P. menispermifolia*; bif= *P. biflora.*

Larval diet also affected size and weight of both species. Forewing size of *H. cydno* (ANOVA, *F*_3,39_= 5.14; *p*= 0.004) was larger and more strongly influenced by larval diet than *H. melpomene* (ANOVA, *F*_3,57_= 4.0; *p*= 0.012) (Figure 2B). *H. cydno* had larger forewings when fed on *P. vitifolia* and *P. biflora*, and smaller on *P. menispermifolia* and *P. platyloba.* In contrast, adults of *H. melpomene* had larger forewings when reared on *P. menispermifolia* and *P. biflora*, and smaller on *P. vitifolia* and *P. platyloba*. Sexual differences in forewing size were not observed in either species (ANOVA, *H melpomene*: *F*_1,59_= 0.369, *p*= 0.546; *H. cydno: F*_1,41_= 1.575, *p*= 0.217). Broadly similar effects were seen for pupal weight, butterfly weight and body size, as for forewing size (Figure S1).

In order to verify whether sequestration versus biosynthesis has a significant effect on fitness of both species, we tested for a correlation between concentration of biosynthesized CNglcs and forewing size (Figure 2C). In the generalist *H. cydno*, even though larval diet strongly affects forewing size, this effect is not correlated with whether they biosynthesize (R^2^= 0.619, *F*_1,41_= 2.707, *p*= 0.108) or sequester (R^2^= 0.09, *F*_1,41_= 4.081, *p*= 0.05) CNglcs. Whilst, in the specialist *H. melpomene,* there is a negative correlation between CNglc biosynthesis and forewing size (R^2^= 0.1339, *F*_1,57_= 8.814, *p*= 0.004), even when its favourite diet was removed of the analyses (R^2^= 0.086, *F*_1,53_= 4.979, *p*= 0.03). This suggests that CNglcs biosynthesis has a fitness cost for *H. melpomene rosina*, which mostly lay eggs on *P. menispermifolia* and sequester CNglcs from it during larval feeding. Additionally, there was a positive correlation between forewing size and concentration of sequestered CNglcs in *H. melpomene* (R^2^= 0.1466, *F*_1,57_= 9.979, *p*= 0.003), indicating that larvae that are better sequestering CNglcs tend to turn into bigger butterflies.

## DISCUSSION

We documented, for the first time, plasticity in CNglc composition and concentration for both *H. melpomene rosina* and *H. cydno chioneus* in response to their larval diet (Figure 1 and 2). We confirmed that when feeding on a plant with cyclopentenyl CNglcs that can be sequestered (*i. e.* deidaclin in *P. menispermifolia*), both butterfly larvae invest less in biosynthesis of aliphatic CNglcs, a trade-off that has previously been proposed at the level of inter-species comparisons [23][16]. This plasticity should facilitate *Heliconius* butterflies adapt to exploit different *Passiflora* hosts, they could utilize plants regardless of their CNglc profile because they can maintain their defences through biosynthesis when sequestration is not possible.

Regardless of how they acquired their cyanogenic defences, both butterflies gained similar total concentration of CNglcs when raised on their natural host range (*P. platyloba, P. menispermifolia* and *P. vitifolia*). A similar pattern has been observed in the six-pot burnet moth *Zygaena filipendulae*, another rare example of lepidopteran that can both *de novo* biosynthesize and sequester their chemical defences [25]. *Z. filipendulae* balance their cyanogenic content with biosynthesis in the absent of sequestration, however with deleterious consequences for their growth[26][27]. It is likely that, as in *Zygaena* moths, *Heliconius* have adaptations to optimize the energetic cost of their toxicity: deactivating the biosynthesis of CNglcs when these compounds are available for sequestration and reactivating it when they are not.

Although adult size and weight of *H. cydno* were strongly influenced by their larval diet (Figure 2), these differences were not correlated with whether they acquired their CNglcs through biosynthesis or sequestration. This suggests that plasticity in the generalist species does not come with a significant energetic cost. In contrast, *H. melpomene* grows bigger (Figure 2B and S1) when favouring sequestration over biosynthesis, suggesting that it has adapted to its specialist lifestyle and has a significant cost to the plasticity involved in switching host plants.

Smiley (1978) emphasized that ecological factors involved in the initial choice of a host plant might not be the same that led to the maintenance of this preference. It seems likely that the Panamanian *H. melpomene* only recently evolved a preference for *P. menispermifolia.* Once this oviposition preference established, selection for digestive adaptations to maximise the larval performance on this diet would take a place - *e. g.* increasing the efficiency of CNglc uptake from *P. menispermifolia* as we observed in this study (Figure 1). Local and recent adaptation to larval feeding on *P. menispermifolia* might also explain why *H. melpomene* performs only slightly better on this diet (Figure 2B and S1). Nonetheless, for the preferred host *P. menispermifolia* we have shown, for the first time, that this is a good host for *H. melpomene*, but a less optimal host for *H. cydno*.

In Panama, avoidance of interspecific competition is likely to be a major force shaping the evolution hostplant range, since coexistent *Heliconius* species rarely shared oviposition preference for the same *Passiflora*: *H. erato* lays eggs preferably on *P. biflora*, *H. hecale* on *P. vitifolia*, *H. sara* on *P. auriculata and H. melpomene* on *P. menispermifolia*[21][28]. Niche partitioning not only happens for *Passiflora* hosts, but also at microhabitat level: whereas most *Heliconius* species, including the comimics *H. melpomene* and *H. erato*, are found in open secondary forest, *H. cydno* and *H. sapho* are typically present in the closed-canopy [29]. A similar pattern of resources partitioning (plant and microhabitat) occurs in Colombia [19]. Thus, interspecific competition might have led *H. melpomene* to evolve specialized oviposition preferences for *P. menispermifolia* and pushed *H. cydno* to inhabit forest where *Passiflora* species are less abundant and a generalist strategy might be favoured. The phenotypic plasticity in their biochemistry enabled *Heliconius* butterflies to widen their range of *Passiflor*a host and led to niche diversification while maintaining their chemical defences, allowing the coexistence of multiple *Heliconius* species.

Finally, the vast majority of aposematic moths and butterflies sequester their toxic compounds from their larval host, emphasizing the importance of this process in the coevolution between plants and lepidopterans [30]. In turn, many *Passiflora* species seems to have modified their cyclopentenyl CNglcs to disable sequestration by heliconiines [16]. Here, we show that the two modified CNglcs passibiflorin (*bis*-glycosilated) and tetraphylli-B sulphate (sulphonated) were not sequestered by both *Heliconius* species, suggesting counter-evolution in the plants to deter their herbivores.

Our findings, based on Heliconius butterflies and its Passiflora host, highlight the importance of phenotypic plasticity in biochemical traits for the diversification of herbivorous insects. A large proportion of global biodiversity is represented by tropical herbivorous insects, so understanding how genetic and plastic traits allow species to adapt their host niche and permit species to coexist is an important step towards understanding biodiversity.

---

**Figure S1.**
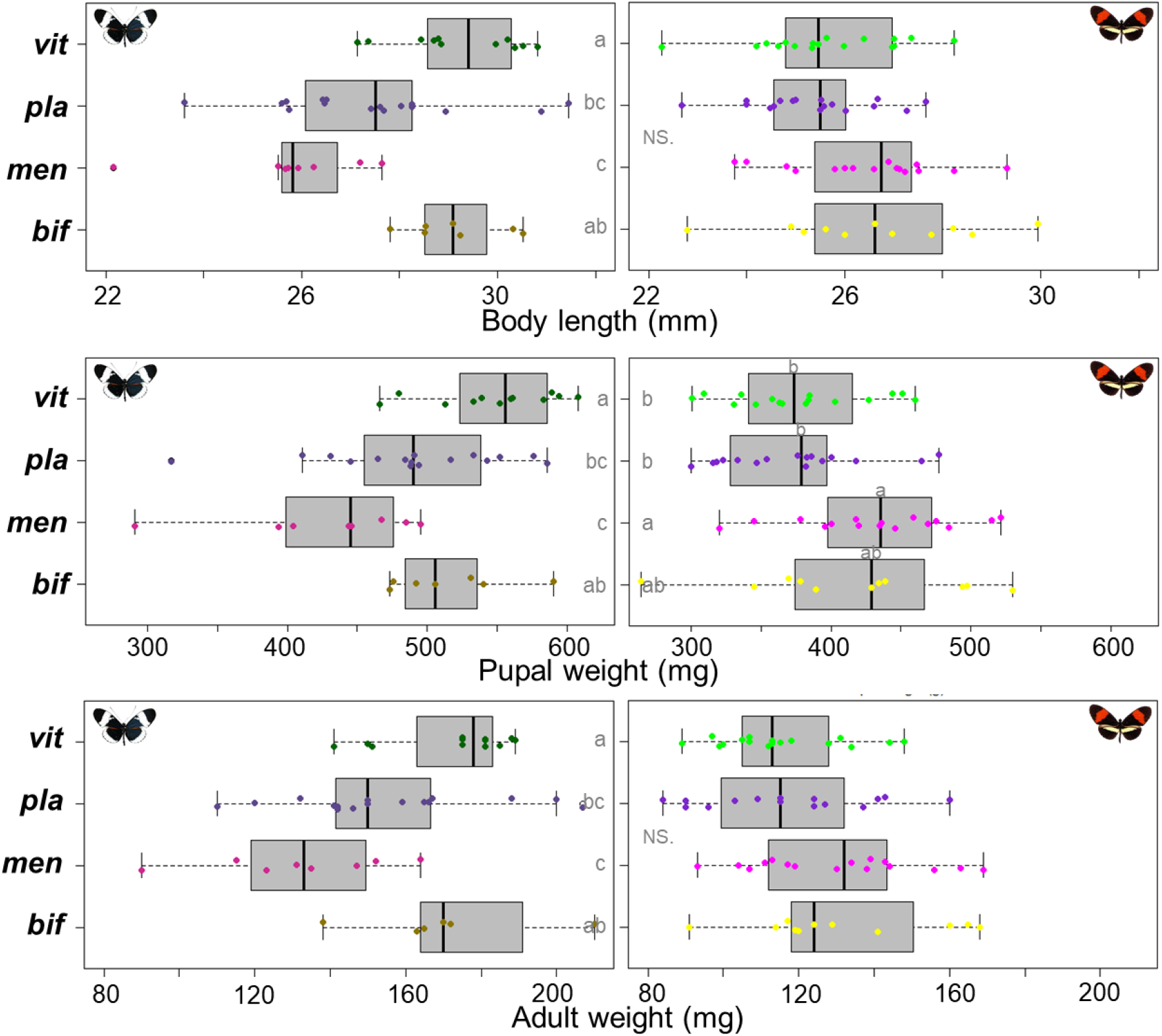
Effect of larval diet on body length, pupal weight and adult weight of *H. cydno* (left) and *H. melpomene* (right). Letters over boxplots correspond to post-hoc comparisons (Tukey HSD) within butterfly species, where different letters indicate statistically significant treatments. Legend: vit= *P. vitifolia,* pla= *P. platyloba,* men= *P. menispermifolia*; bif= *P. biflora.*

